# The perception of auditory motion in sighted and early blind individuals

**DOI:** 10.1101/2022.09.11.507447

**Authors:** Woon Ju Park, Ione Fine

## Abstract

Motion detection is a fundamental property of the visual system that plays an important role across many of the human senses. In vision, motion processing is classically described using a motion energy model which assumes spatiotemporally selective (i.e., non-separable) filters that capture the smooth continuous change in spatial position over time afforded by moving objects. However, in the case of audition, it is still not clear whether or not the primary mechanisms underlying motion sensitivity are spatiotemporally selective. We used a psychophysical reverse correlation paradigm, where participants discriminated the direction of a motion signal in the presence of spatiotemporal noise, to determine whether the filters underlying auditory motion discrimination were spatiotemporally separable or non-separable. We then examined whether these auditory motion filters were altered as a result of early blindness. We found that both sighted and early blind individuals have separable filters. However, early blind individuals show increased sensitivity to auditory motion, with reduced susceptibility to noise, with filters that were more accurate in detecting motion onsets/offsets. An ideal observer model suggested that the reliance on separable filters may be more optimal given the limited spatial resolution in auditory input.

## INTRODUCTION

Understanding the motion of objects in the environment is a fundamental task in sensory processing. In vision, it is well-known that the earliest stage of specialized motion processing (area MT) contains neurons whose tuning is spatiotemporally non-separable, specialized for the detection of continuous, object motion in visual space. In contrast, the question of how the human brain processes auditory motion has been debated for over three decades. Specifically, it is not clear whether auditory motion perception is inferred from “snapshots” of individual static sounds at different locations, Figure 1A, or relies on dedicated non-separable “motion detectors” as in vision, Figure 1B.

**Figure 1.**
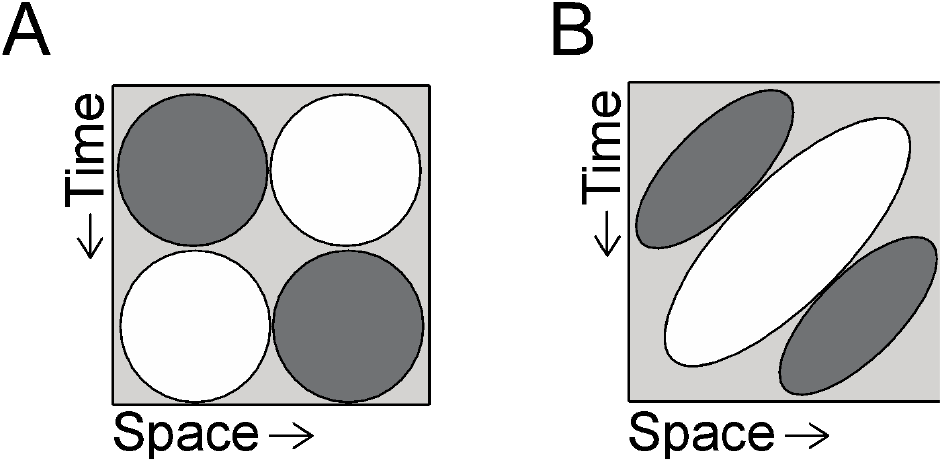
A. Space–time tuning. A. Separable spatial-temporal tuning can be described as the outer product of separate 1D filters in space and time. B. In contrast, for motion selective non-separable filters, the contour of the receptive field is tilted with respect to the coordinate axes. The maximal activation of the filter occurs for stimuli whose location in space in time can be described as v = x/t, where v is velocity, t is time and x is spatial location.

Previous human psychoacoustical findings have been inconclusive. Some studies have measured the motion aftereffect (MAE), based on the longstanding notion that selective adaptation requires the existence of specialized mechanisms tuned for the adapted sensory feature. While adaptation to auditory motion can produce a measurable MAE, the effect is highly stimulus specific (e.g., does not generalize across frequencies) (Dong et al., 2000; Grantham, 1989; Grantham & Wightman, 1979) and is much weaker (Neelon & Jenison, 2003) than the adaptation observed using visual motion stimuli (Anstis et al., 1998). In addition, auditory speed judgments tend to be dominated by duration and distance cues, a finding more consistent with separable tuning (Freeman et al., 2014; see Carlile & Leung, 2016 for a review).

The neurophysiological evidence is similarly inconclusive. Some animal studies do find direction-selective cells (Ahissar et al., 1992; Altman, 1968; H. Jiang et al., 2000), but their selectivity seems to result from selectivity to acoustic features associated with motion stimuli, rather than showing selectivity to motion per se (Grzeschik et al., 2010; Ingham et al., 2001; McAlpine et al., 2000). fMRI studies in humans have consistently shown greater responses to auditory motion as compared to static stimuli in the right planum temporale (rPT; Alink et al., 2012; Baumgart et al., 1999; Ducommun et al., 2004; Poirier et al., 2005; Warren et al., 2002). However, the responses in this region to auditory motion are comparable to static stimuli that randomly change location over time (Smith et al., 2004, 2007), suggesting that PT likely encodes changes in sound position rather than genuine auditory motion.

We used a psychophysical reverse correlation paradigm to estimate the shape of the “filters” for discriminating auditory motion in human listeners with neurotypical visual and auditory histories. Psychophysical reverse correlation paradigms have been used to characterize the mechanisms underlying a wide variety of visual tasks (Abbey & Eckstein, 2002; Ahumada, 1996; Murray, 2011; Neri et al., 1999), and the “perceptive” field properties estimated from these paradigms have shown remarkable similarities to receptive fields of single neurons (Neri & Levi, 2006). Our particular paradigm is a direct analogue of a previous study that observed clear spatiotemporal non-separability in the perceptual filters estimated for discriminating visual motion direction (Neri, 2014), which closely resembled the known spatiotemporal selectivity of neurons within area MT.

Participants discriminated the direction of motion of a near-threshold auditory motion signal embedded in bursts of static auditory noise randomly presented over space and time. By characterizing how the spatiotemporal structure of the auditory noise systematically influences the task decision, we aimed to extract the spatiotemporal features that contribute to the perception of auditory motion direction.

We then used the same reverse correlation paradigm in individuals who became blind early in life. Early blind individuals rely heavily on auditory information for their everyday navigation. Understanding object motion (e.g., cars) based entirely on auditory information is an ecologically critical task that blind individuals are forced to learn. One previous study did show enhanced auditory motion selectivity as a function of early blindness (F. Jiang et al., 2014), however this was a highly complex task requiring detecting the overall direction of motion of multiple incoherent moving sound sources. A second recent study, examining speed processing, found that early blind individuals were, if anything, less sensitive to speed, and relied predominantly on stimulus duration when making speed judgments (Bertonati et al., 2021). Here we examine whether the ‘perceptive’ filters underlying motion discrimination are altered by early blindness.

## METHODS

This study was approved by the University of Washington’s Institutional Review Board and carried out in accordance with the Code of Ethics of the Declaration of Helsinki. Informed written consent was obtained from all participants prior to conducting the experiments.

### Participants

Adults were recruited from the greater Seattle, WA community. Participants included 8 visually typical observers and 8 early blind individuals. The two groups were matched for age and music experience (see Tables 1 & 2). All participants reported normal hearing and no history of psychiatric illness.

**Table 1.** Visually typical participants’ characteristics.

|  | Sex | Age | Music Training<br>(yrs) |
| --- | --- | --- | --- |
| Sighted1 | F | 62 | 55 |
| Sighted2 | M | 35 | 28 |
| Sighted3 | M | 34 | 0 |
| Sighted4 | F | 62 | 5 |
| Sighted5 | M | 73 | 66 |
| Sighted6 | M | 64 | 50 |
| Sighted7 | M | 61 | 45 |
| Sighted8 | M | 47 | 0 |

**Table 2.** Early blind (EB) participants’ characteristics.

|  | Sex | Age | Music<br>Training (yrs) | Blindness<br>Onset | Cause of Blindness | Light<br>Perception |
| --- | --- | --- | --- | --- | --- | --- |
| EB1 | M | 55 | 50 | Born blind | Congenital glaucoma | No |
| EB2 | M | 28 | 20 | Age 2 | Retinoblastoma | No |
| EB3 | M | 36 | 0 | Born blind | Leber's congenital<br>amaurosis | No |
| EB4 | F | 59 | 0 | Born blind | Retinopathy of prematurity | No |
| EB5 | M | 69 | 50 | Born blind | Retinopathy of prematurity | No |
| EB6 | F | 58 | 50 | Age 2 | Optic atrophy | No |
| EB7 | F | 68 | 30 | Born blind | Retinopathy of prematurity | Low |
| EB8 | M | 50 | 1 | Born blind | Optic nerve hypoplasia | Low |

### Stimulus

Auditory stimuli were delivered through Etymotic ER-2 insert earphones at a sampling rate of 44,000 Hz. The 3D auditory space was simulated based on a basic physics model, using interaural level and time differences, and the decrease in volume as a function of the distance to the observer (Doppler shift and head shadow were not modeled). The stimuli were generated and presented using MATLAB and Psychtoolbox (Brainard, 1997).

#### The stimuli consisted of two components

*signal motion* and *background noise bursts*. Both stimuli were broadband noise, created by generating Gaussian noise in the time domain, which was then bandpass filtered between 500-14,000 Hz in the Fourier domain (fast Fourier transform) and was projected back to the time domain (inverse fast Fourier transform).

The signal motion continuously traveled either leftward or rightward (Figure 2A) on a given trial. We simulated a constant-velocity stimulus traveling from +/-15m from the observer, along a frontoparallel direction (a straight-line oriented perpendicular to the listener’s facing direction) at a distance of 0.8 m, centered at the midline. The signal motion lasted 500 ms and had a linearly ramped onset and offset of 50 ms. The volume of the signal motion was adjusted throughout the experiment using two interleaved two-down one-up staircases which held performance accuracy at approximately 65%.

**Figure 2.**
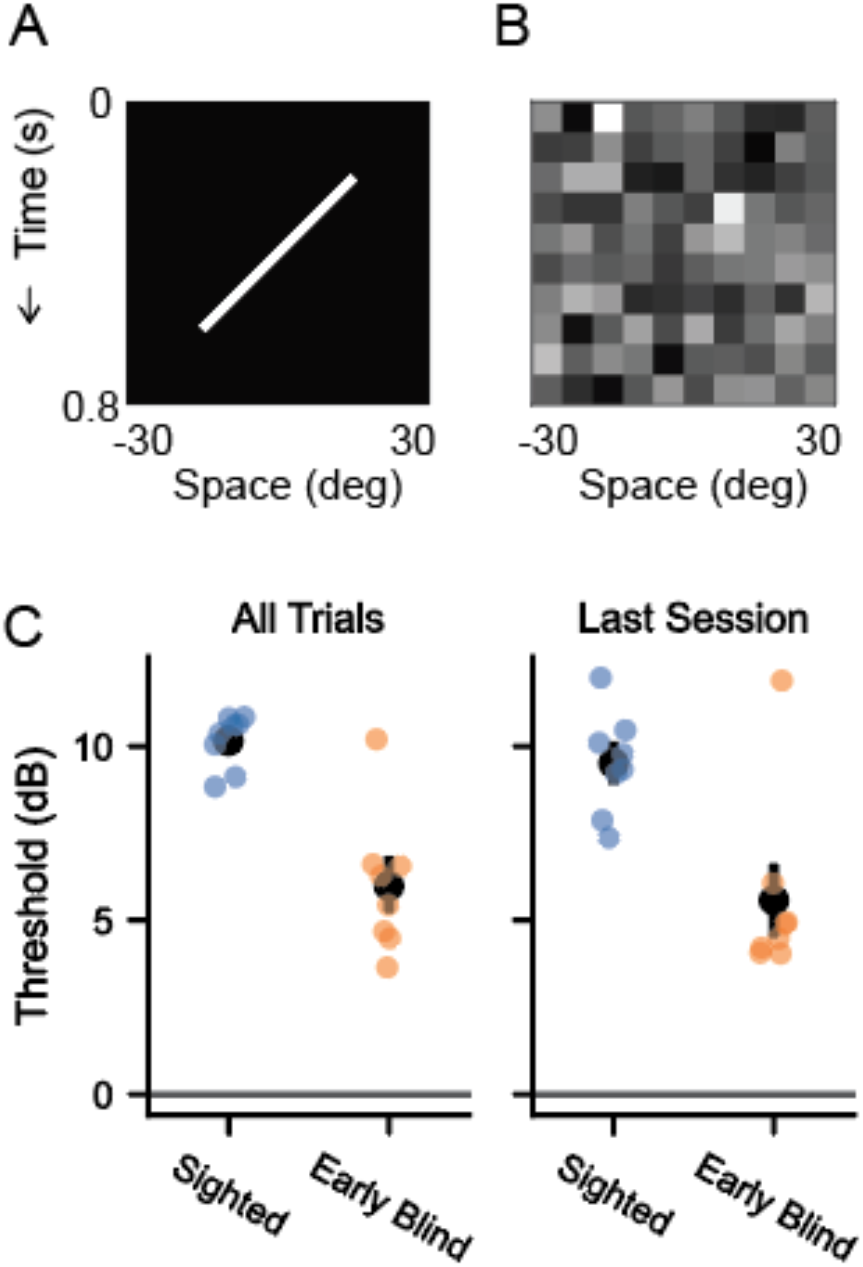
Auditory motion filters estimated using a psychophysical reverse noise technique that was a direct analogue to Neri et al., 2014. A. Participants were asked to discriminate the direction of signal motion (broad band noise, left vs right, leftward motion depicted in the panel). B. Spatiotemporal noise was added to the moving signal. Intensity in each cell in A and B represents sound amplitude (scale = [0 1], from black to white). C. Early blind individuals can identify the direction of motion at lower levels of signal amplitude than sighted individuals, left panel shows thresholds based on the data from all six sessions (1000 trials/session), right panel shows thresholds for the 6th (final) session. Individual thresholds (colored circles) and group means (black circles) with single standard error bars are shown.

The background noise bursts were discrete sounds simulated on a 10 × 10 grid (Figure 2B) that spanned space (-/+30 °) and time (0-800 ms). At each moment in time, there were 10 simultaneous noise bursts (of varied amplitude), one for each location in space. The amplitude of the noise bursts at each spatiotemporal location within the space-time grid was randomly and independently selected from a Gaussian distribution, resulting in an approximate range of 5-49 dB (mean = 39 dB).

### Task

On each trial the observers reported whether the auditory motion stimulus was moving leftward or rightward via a button press and were given auditory feedback (a brief beep if they were correct).

At the end of each block (200 trials) there was a brief rest period, after a mandatory break of 30s participants could press any key to begin the next block of trials. Each session contained 1000 trials, and each participant carried out 6 sessions. The first block was discarded from the analyses to limit early practice effects.

### Analysis

#### Derivation of Signal Motion Thresholds

Each participant’s threshold to discriminate the signal motion was estimated by fitting a Weibull function using the psignifit toolbox (Schütt et al., 2016) to all trials from each staircase within a session. This resulted in six threshold measurements, which were then averaged.

#### Derivation of Spatiotemporal Perceptual Filters

The perceptual filters for hearing auditory motion were estimated for each participant, *solely* using the characteristics of noise bursts, independent from the changes in signal motion amplitude. The filters were derived based on Neri (2014), who used an analogous paradigm for estimating visual motion filters in visually typical observers. Here, *N*^[*q,z*]^ denotes the noise sample over space (*x*) and time (*t*), and ⟨ ⟩ represents the average across trials. Filters were constructed by sorting trials into four categories based on the direction of the signal direction (left (*q* = 1) or right (*q* = 0)) and participants’ response (correct (*z* = 1) or not (*z* = 0)). The resulting filter (*F*) is derived:

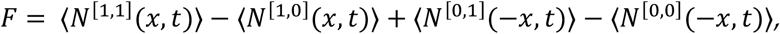

where the spatial axis is mirror-inverted (− *x*) for noise samples on trials with rightward-signals, to align the filter orientations.

#### Comparison of Separable and Non-separable Models

We assessed whether human listeners use spatiotemporally separable or non-separable models for hearing auditory motion in two ways.

We began by comparing two simple models which differed in their spatiotemporal separability (Figure 1B). The separable filter assumed Gaussian tuning in both space and time, described with three free parameters: amplitude, center, and sigma. The center locations for spatial and temporal tuning were assumed to be symmetric. Four identical Gaussians were created based on these parameters, one in each of the four quadrants in the space-time grid. For the leftward tuned filter, the amplitude of the top-left and bottom-right Gaussians was positive, and the amplitude of the top-right and bottom-left Gaussians was negative. The rightward tuned filter was constructed as the reciprocal combination of positive and negative Gaussians.

The non-separable filter was modeled as an oriented Gaussian. Once again there were three parameters: amplitude, sigma, and orientation. To create inhibitory responses, filter values were scaled between [−1, 1]. *amplitude*.

For each individual, we found the parameter values that minimized the mean square distance between the model filter and the measured perceptual filter for that individual, using both the separable and the non-separable models. Goodness of fit was assessed by taking the dot product of the predicted and the measured perceptual filters.

Next, we examined spatiotemporal separability using a model that could directly characterize how “separable” the filters are in space and time. Specifically, two separate surfaces (*I*_1_, *I*_2_) were defined (Figure 5A), each defined as the outer product of a grating in space and a grating in time:

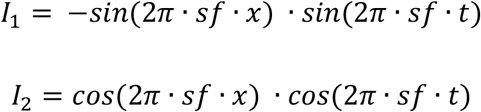

where *sf* is spatial frequency. (An additional free parameter, *θ*, was used to find the best fitting grating orientation).

The linear combination of these two surfaces, *M* = *I*_1_ + *a*. *I*_2_, ranges between a separable surface (*I*_1_) when *a* = 0 and a non-separable oriented grating when *a* = 1, as shown in Figure 5B.

Thus, the parameter *a* provides a measure of spatiotemporal separability that can be fit to each participant’s measured perceptual filters.

#### Characterizing Effects of Experience on Auditory Motion Filters

To examine how early blindness alters auditory motion filters, we first characterized how the sighted and early blind individuals weighted the spatiotemporal properties of the noise differently. To do this, we ran 1,000 Monte Carlo simulations assuming no group differences in the obtained perceptual filters by randomly assigning a group (sighted or early blind) to each participant’s filter while maintaining the number of participants identical to the study. We then compared this ‘null’ difference filter distribution to the actual difference between the averaged sighted and early blind filters. The cells within the space x time grid that had values outside of 95% percentile (one-sided) were considered significantly different between the two groups.

Second, we fit a separable model that assumes Gaussian tuning in both space and time to the measured perceptual filters. The model was similar to the separable model mentioned above but was expanded by including additional free parameters that allowed us to characterize the changes in spatiotemporal tuning in more detail. Specifically, the model had had five free parameters: amplitude, center, and width for spatial tuning, as well as center, and width for temporal tuning. The parameters were estimated for each participant and were statistically compared between the groups.

We also tested whether the measured auditory motion filters predict behavioral performance. We correlated the estimated parameters from the expanded separable model to participants’ independently measured perceptual thresholds for discriminating signal motion. The correlation analyses were performed separately for each group.

## RESULTS

### Early blind individuals can discriminate auditory motion better than sighted

Figure 2C shows the signal motion amplitude that resulted in 65% accuracy, for both sighted and early blind individuals. Overall, sighted individuals required signal motion that was approximately three times louder than the mean background noise level. In contrast, early blind individuals were able to perform the same task with significantly lower signal amplitude (t(14) = -5.47, p < .001), with thresholds that were almost 50% lower than sighted individuals. This enhanced performance was still apparent when the data from only the last session were used to estimate the thresholds (t(14) = -3.71, p = .002). The perceptual advantages accrued from auditory experience due to blindness are still apparent even after both participant groups had experienced 5,000 trials of practice.

### Both sighted and early blind individuals have separable auditory motion filters

Figure 3A shows estimated perceptual auditory motion filters for individual sighted and early blind participants. The group average is depicted in Figure 3B. It is worth noting that this measure is independent of the signal motion threshold: the filters were constructed entirely based on the properties of the external background noise, whereas the perceptual threshold was obtained based on the amplitude of signal motion.

**Figure 3.**
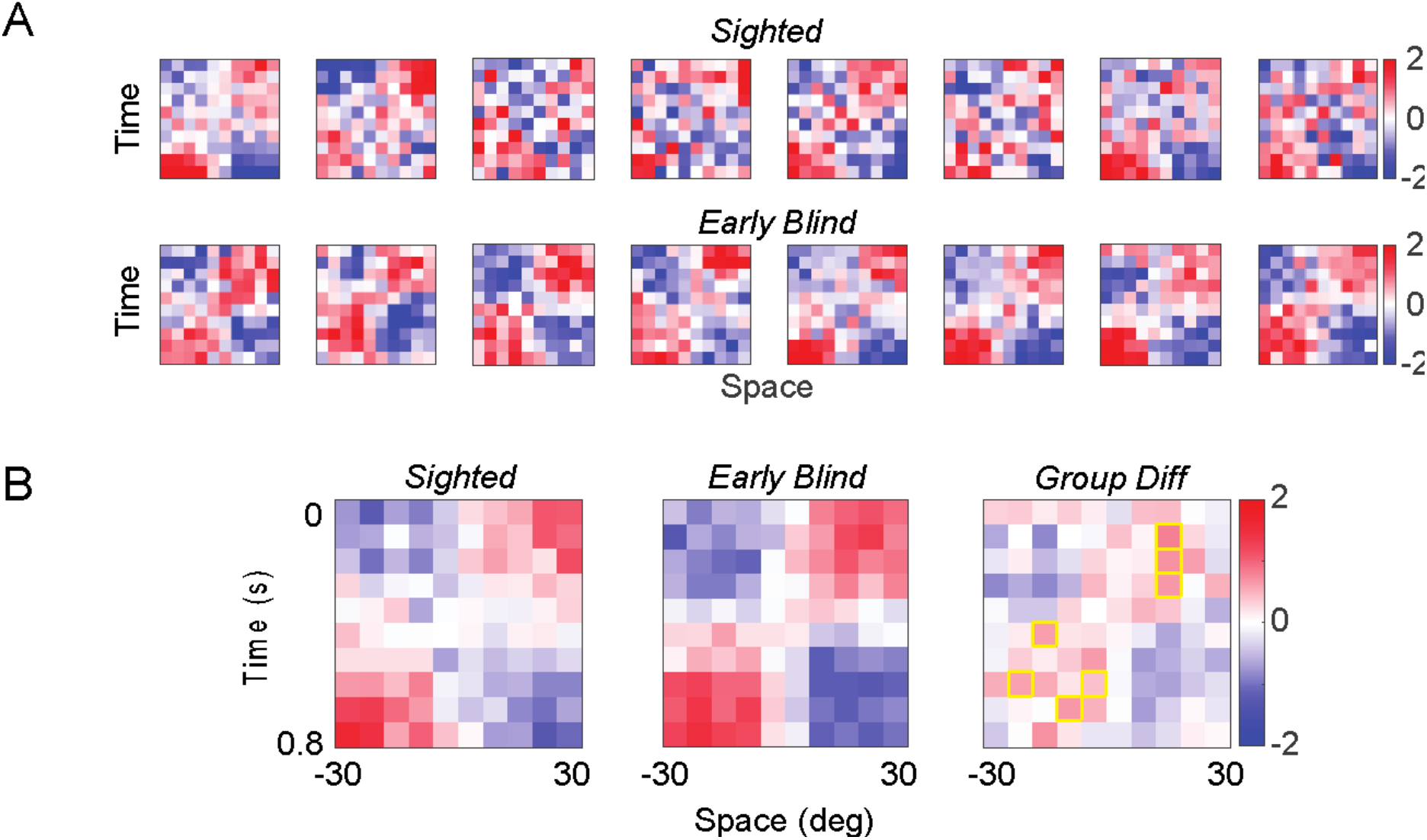
A. Estimated perceptual auditory motion filters for sighted (top row) and early blind (bottom row) individuals. B. Group averaged filters for sighted (left) and early blind (center) individuals. The panel on the right shows the group difference, with yellow outlined cells indicating those that show a significant group difference. The intensity in each cell represents the Z score (measuring the influence of noise at that temporal and spatial location on motion judgments).

Consistent with the “snapshot” model of auditory motion processing, on visual inspection, sighted and early blind individuals both appear to have separable filters. To provide a metric quantifying the separability in the filters, we compared two simple models: one that extracts the sound locations at the onset and the offset (separable model, Figure 4A left) and the other that can continuously track auditory motion (non-separable model, Figure 4A right). The models were separately fitted to each participant’s filter to examine which model explains the shape of the perceptually measured filters better.

**Figure 4.**
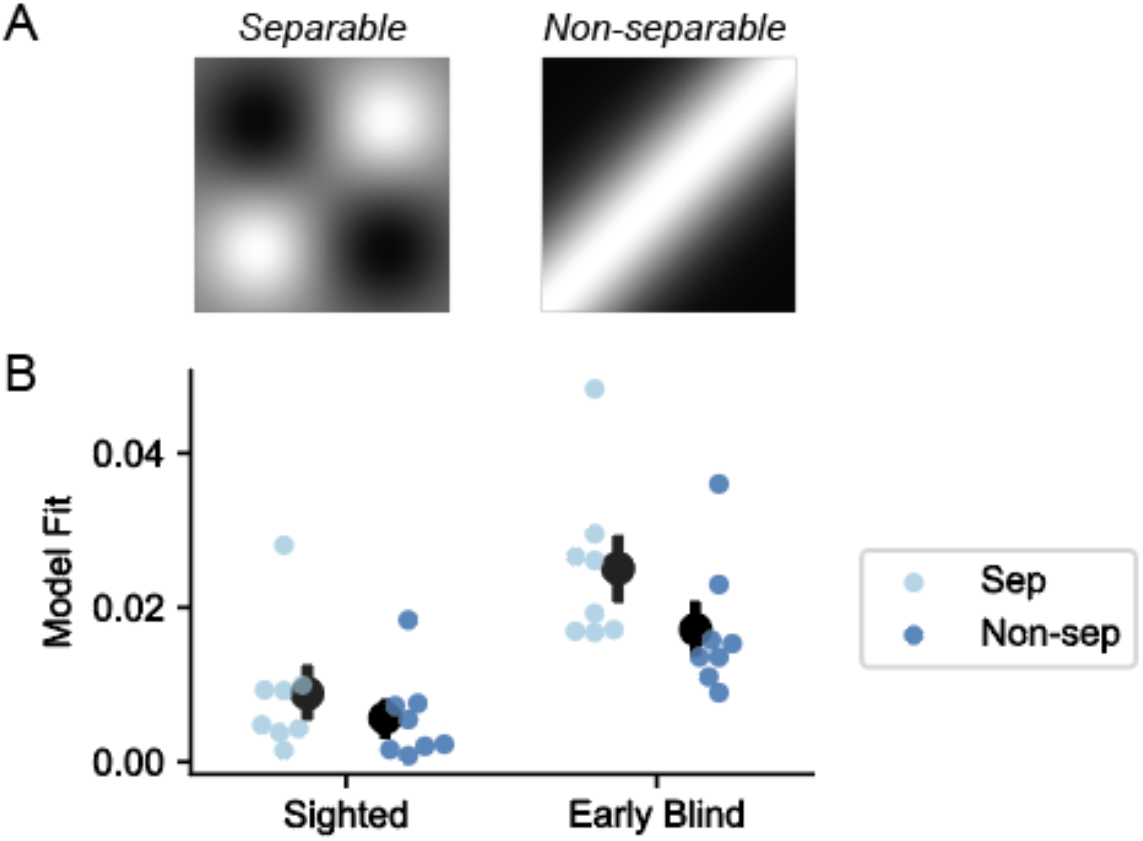
A. Separable (left) and non-separable (right) models fit to individual perceptual filters shown in Figure 3A. B. Model fit illustrating how well the separable vs. non-separable models describe the measured filters. Model fit was assessed by the dot product of the fitted model and measured perceptual filters for each participant (colored circles). Black circles show the group mean. Error bars are standard error of the mean.

As can be seen from Figure 4B, the separable model provided a better fit than the non-separable model (F(1,14) = 56.35, p < .001). Overall, both the separable and non-separable models explained the perceptually measured filters better in early blind individuals (F(1,14) = 10.84, p = .005), and the difference in performance between the two models was greater in early blind than sighted (F(1,14) = 10.24, p = .006) but was significant in both groups (sighted: p = .014; early blind: p < .001).

A separate analysis that directly quantified the “separability” of the perceptual filters similarly suggested that the filters were spatiotemporally separable in both sighted and early blind groups. Figure 5C shows the estimates of the separability parameter (a) for all participants are clustered around 0, rather than 1, implying clear spatiotemporal separability. The fact that the filters were separable in both groups also suggests that the influence of auditory experience on the *shape* of the auditory motion filters is minimal.

**Figure 5.**
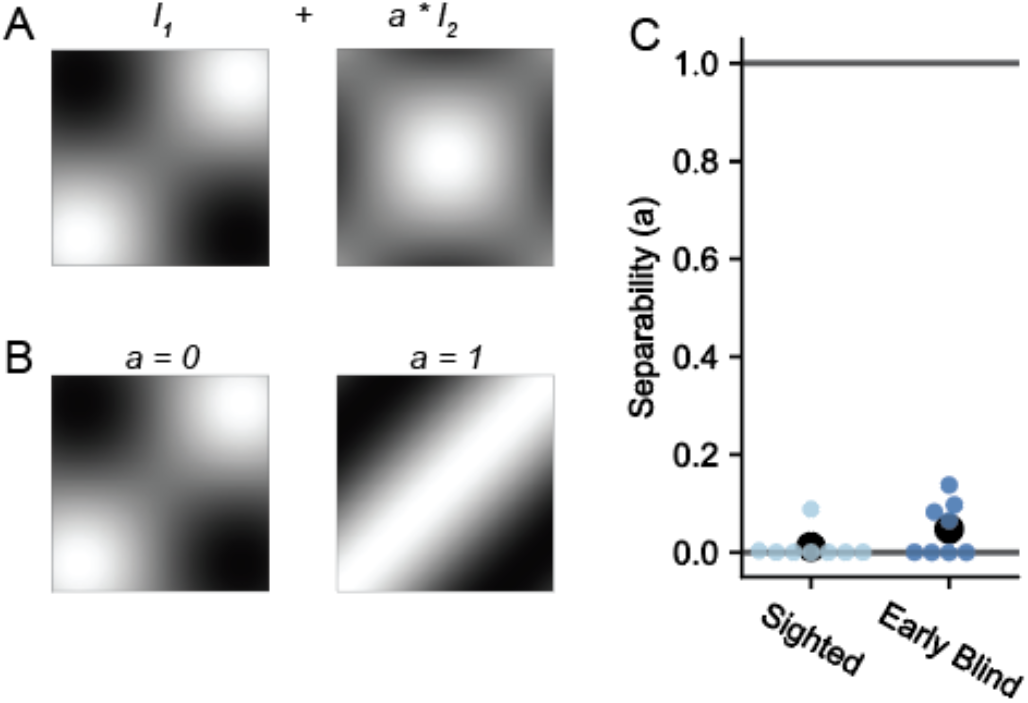
A. Quantification of spatiotemporal separability in the measured auditory motion filters (see Methods for details). B. The parameter *a* characterizes the separability of the individual filters, ranging from separable (0) to non-separable (1). C. Estimated separability parameter (*a*) for each participant (colored circles). Black circles show group mean. Error bars are standard error of the mean.

### Early blindness refines auditory motion filters

Can the enhanced performance in early blind individuals be explained by changes in auditory motion filters? Figure 3C (right) shows the group difference in the measured perceptual filters, showing the regions in space and time which were weighted differently in each group. The colored cells show the regions that were significantly differently between the two groups, as determined by Monte Carlo simulations (see Methods). As can be seen, early blind individuals had their peak filter regions shifted towards the actual signal onset and offset compared to the sighted group.

Parameter estimates from the separable Gaussian model (with five free parameters) showed consistent and significant differences in spatiotemporal tuning in early blind individuals (Figure 6A). Specifically, we observed a significant difference in the center parameters for space (t(14) = -2.4, p = .031), where the center of the Gaussian tuning was more shifted towards the center of the space x time grid in early blind individuals. The group difference in the center parameter for temporal tuning was marginally significant (t(14) = 1.93, p = .074). Consequently, the peak coordinates within the filters constructed using the estimated parameters were more aligned with the onset of the *signal motion* for early blind individuals, while those for the sighted were closer to the start of the background noise (Figure 6B). These results suggest that early blind individuals’ filters enabled more accurate detection of the spatiotemporal onset and offset of the signal motion The amplitude parameter from the model was also larger in early blind than sighted individuals (t(14) = 3.72, p = .002), consistent with the idea that early blind individuals have either larger gain or lower internal noise in their auditory motion processing. The spatial and temporal width parameters did not show a significant group difference (spatial width: t(14) = 1.19, p = .26; temporal width: t(14) = 1.39, p = .19).

**Figure 6.**
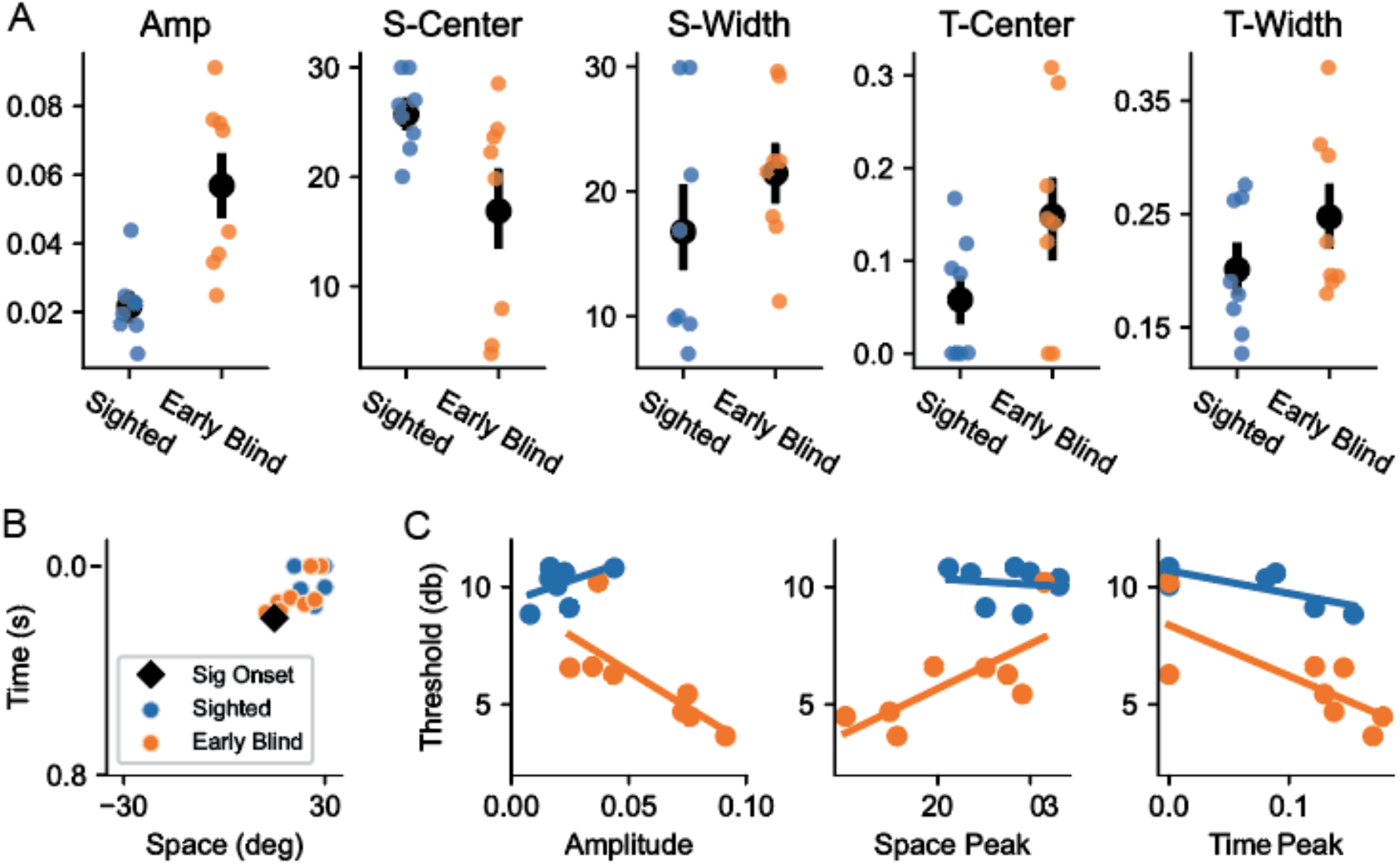
A. Parameter estimates from the separable Gaussian model that had five free parameters. Compared to the sighted group, early blind individuals had filters that had greater amplitude and center tuning more shifted towards center of the space x time grid. B. The peak coordinates within the fitted filters for sighted and early blind individuals. Early blind individuals’ tuning was closer to the actual signal onset (black diamond). C. Results from the correlation analyses examining the relationship between estimated filter properties for sighted (blue) and early blind (orange) individuals. Lines show the linear regression fit.

To examine if the estimated filters predict behavioral performance, we correlated the filter properties with the measured perceptual thresholds (Figure 6C). The results showed that, in early blind individuals, the spatial and temporal peak coordinates within the filter were significantly correlated with perceptual thresholds (space: r(6) = .74, p = .038; time: r(6) = -.76, p = .028), where closer peak to the signal onset was associated with better performance. We also found a significant correlation between amplitude and thresholds in early blind individuals (r(6) = -.75, p = .03), with higher filter amplitude predicting better performance. These results suggest that the shift in spatiotemporal tuning in early blind individuals’ filters explains their improved behavioral performance. For sighted individuals, the peak coordinate on the temporal axis was significantly related to thresholds (r(6) = -.8, p = .018), again showing that the estimated filter properties are related to individuals’ auditory motion perception. None of the other correlation analyses showed a significant relationship (all p’s > .25).

### Separable filters may be a consequence of broad auditory spatial tuning

The results thus far demonstrate that auditory motion discrimination likely relies on spatiotemporally separable filters, which can be refined based on extensive auditory experience. Why is it that visual direction discrimination relies on non-separable filters, while that same task, done in the auditory domain relies on separable filters? One major difference between the two modalities is that visual spatial tuning is exquisitely precise (individuals can detect differences of the location of less than a degree) whereas auditory spatial tuning is quite coarse (sensitive to shift in spatial location of ∼2-3 degrees) (Alais & Burr, 2004), see Discussion. To examine whether this difference in spatial tuning width accounts for the shift from non-separable to separable tuning, we carried out an ideal observer analysis, comparing the performance of separable and non-separable filters on our experimental task as a function of tuning width (Figure 7).

**Figure 7.**
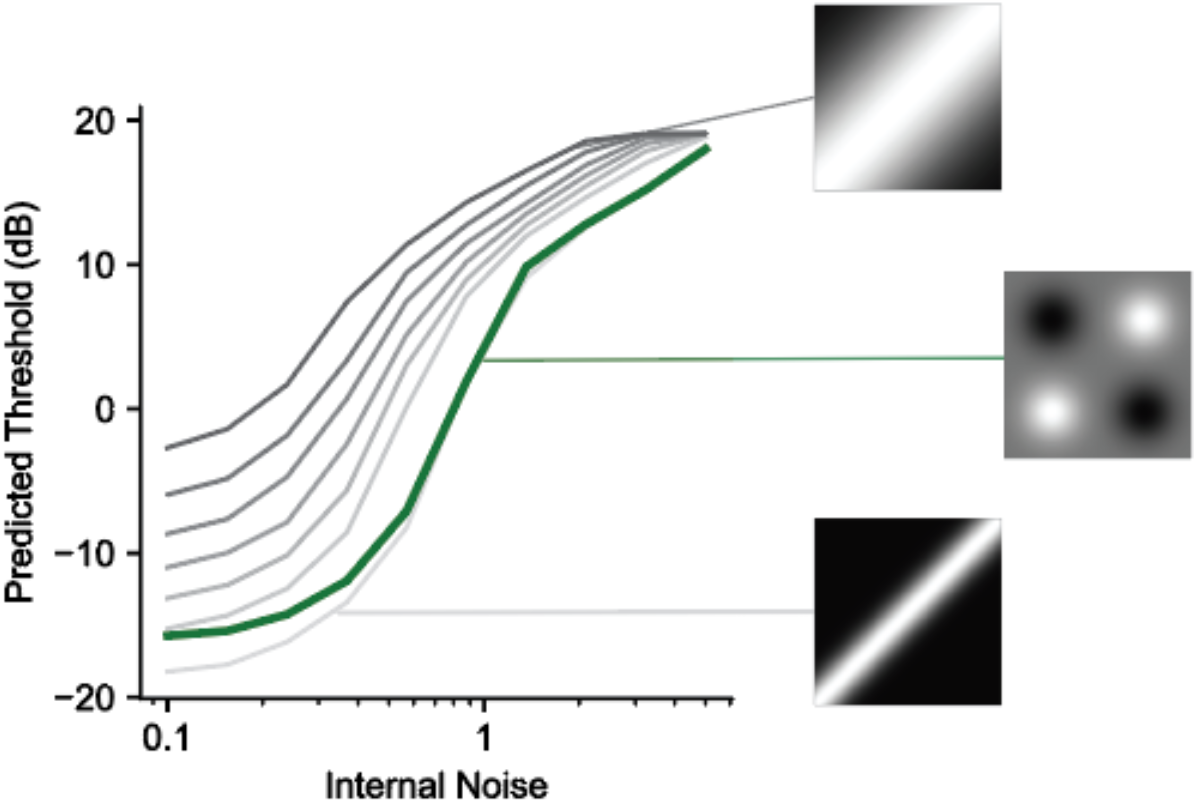
Performance of an ideal observer model simulating a set of non-separable models with varying tuning width from narrow (lighter grey) to broad (darker grey) as well as a separable model constructed using the average spatial tuning from our psychophysical data (green). Separable tuning appears to be optimal except under conditions of very low internal noise and very narrow spatial tuning.

We simulated seven non-separable models with varying tuning widths and one separable model, with Gaussians perfectly aligned with the signal spatial and temporal onset/offset and a sigma width that was the average of the spatial width parameter across all participants. The amplitude of the models was fixed at 1.

The response, *R*, of each filter, *F*, to each stimulus, *S*, was calculated as the dot product of the stimulus and the filter with added zero-mean normally distributed internal noise, *N*, with standard deviation proportional to the square root of ⟨*S* ∗ *F*⟩ (the mean filter response).

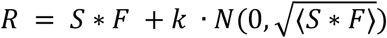

We assumed a correct response when the appropriately oriented filter (e.g., leftward) had a larger response than the opposite (e.g., rightward) tuned filter. The amplitude of the signal motion was varied to find the perceptual threshold as a function of varying internal noise. Internal noise level (*k*) was simulated at 10 levels ranging between 0.1 and 5. The filters were normalized such that the sum of the responses within the excitatory and inhibitory regions are 1 and -1, respectively.

The results show a clear advantage of having a separable filter when there is broad spatial tuning is assumed. Among the non-separable models (grey curves), the most narrowly tuned model performed the best across all internal noise levels simulated. Remarkably, the separable model (green curve) performed as well as the most narrowly tuned non-separable model under almost all noise levels. The only condition in which the separable model did not outperform the non-separable models was with the most narrowly tuned model at very low amounts of internal noise. Thus, spatiotemporally separable filters may be a more optimal solution when the spatial resolution of the incoming sensory information is limited.

## DISCUSSION

Here we used a psychophysical reverse correlation paradigm, where participants discriminated the direction of a motion signal in the presence of spatiotemporal noise, to examine the filters underlying auditory motion discrimination in early blind and sighted individuals.

### Auditory motion perception is mediated by separable filters

We found, for both sighted and early blind individuals, that auditory motion discrimination was mediated by separable filters. This contrasts with what has been observed using the visual analog our auditory task (Neri, 2014), which showed reliance on spatiotemporally non-separable filters for perceiving for visual motion. Thus, direction discrimination is carried out very differently across auditory and visual modalities.

This difference in tuning between auditory and visual motion perception is likely driven by the statistics of the available sensory input. In vision, it is well-established that the resolution for encoding space is exceptionally good: neurons directly represent each point in retinal space. Humans can detect visual spatial offsets as small as 2 to 5 arcseconds (Westheimer & Levi, 1987; Westheimer & Mckee, 1977), and thresholds for detecting visual displacements of successively presented static visual stimuli can be as low as 5 min (MacVeigh et al., 1991; Verdon-Roe et al., 2006). In contrast, in audition, space must be inferred by a combination of cues, including interaural time and level differences. Auditory spatial tuning is broad, within both the primary auditory cortex and PT (Derey et al., 2016; van der Heijden et al., 2018) and neural tuning to sound-source locations seems to be represented by an opponent process, based on differences in the activity of two broadly tuned channels formed by contra- and ipsilaterally preferring neurons (Derey et al., 2016; McAlpine et al., 2001; Stecker et al., 2005). In addition, free-field localization of spectrally rich auditory stimuli produces discrimination thresholds much larger than those of vision, on the order of 1° (Mills, 1958; Perrott & Saberi, 1990).

Using an ideal observer analysis, we found that this difference in spatial resolution between visual and auditory modalities may explain why non-separable filters mediate performance in visual tasks whereas separable tuning mediates performance in auditory tasks. Non-separable filters are only optimal for direction discrimination if tuning is narrow and there are low amounts of internal noise, conditions more likely to occur for visual than auditory motion stimuli.

### Auditory motion processing is enhanced in early blind individuals

Our results also demonstrate how the spatiotemporal tuning of auditory motion filters are altered by early blindness. Early blind individuals showed increased sensitivity to auditory motion, with significantly lower target motion amplitudes required to discriminate the direction of motion at 65% accuracy. This enhanced performance could successfully be predicted from their measured filter properties: early blind filters had larger amplitudes and were more accurate in detecting motion onsets/offsets.

The larger amplitude of the filters of early blind individuals can be interpreted in two ways. One possibility is that this parameter reflects neural mechanisms that are more narrowly and ‘cleanly’ tuned. Previous studies in vision literature suggest that broadly tuned templates can lead to decreased efficiency in filtering out external noise (Dosher & Lu, 1998; Lu & Dosher, 2004), and the use of less well-tuned filters can result in noisier behavioral responses (Beck et al., 2012). Thus, the more refined tuning of the early blind individuals’ filters may have additionally manifested itself as increased amplitude in the filter estimates. Alternatively, the larger amplitudes might reflect a reduction in internal noise (occurring either before or after the filter itself), which would effectively increase the gain in neural responses to the signal. In our experiment, it was impossible to distinguish between these two accounts, but future work, varying the level of external noise, could be used to differentiate these two models.

The neural basis of the refinement of auditory motion filters in early blind individuals is still unclear. One possibility is that it is mediated by extensive auditory experience. Audition is the only source of sensory input that can provide information about distant space other than vision. Thus, early blind individuals must rely heavily on auditory information to navigate and understand where objects are moving in the environment, which might result in experience-based plasticity in their auditory motion processing. There is analogous evidence that musicians also have enhanced basic auditory abilities (Sanju & Kumar, 2016). However, it is not clear whether these results reflect extensive training or differences in innate ability. Our findings certainly suggest that the differences we see in early blind individuals require *extensive* experience —in our study, there was little change in either early blind or sighted individuals’ performance over 6,000 trials (∼6 hours) of practice.

A second possibility is that the alterations in auditory motion processing we observed are driven by sensory reorganization that takes place due to visual deprivation, including cross-modal plasticity in the brain. There is now a large amount of evidence suggesting that visual motion area hMT+ responds to auditory motion in early blind individuals (Bedny et al., 2010; Dormal et al., 2016; Lewis et al., 2010; Poirier et al., 2006) and this region may even ‘co-opt’ the function of rPT (Dormal et al., 2016; Jiang et al., 2014, 2016), a region associated with auditory motion processing in sighted individuals. This raises the possibility that the enhanced spatiotemporal tuning in auditory motion filters of early blind individuals may be mediated by recruitment of deprived visual motion area, hMT+.

The finding of separable filters in early blind individuals, in this context, is somewhat surprising. Given that hMT+ has been shown to have non-separable spatiotemporal filters for processing visual motion, then *if* auditory motion is processed within hMT+ in early blind individuals, then this must result in a shift from non-separable to separable templates—a significant modification of hMT+ normal computational operations.

Alternatively, it is possible that the auditory motion responses observed in hMT+ reflect spatial and temporal signals from either subcortical (perhaps cross-modally recruited superior colliculus (Coullon et al., 2015), or cortical (Gurtubay-Antolin et al., 2021) areas. If so, the ability to classify direction of auditory motion in MT might be mediated by properties other than spatiotemporal tuning, such as retinotopic organization. It remains possible that these signals in hMT+ nonetheless play an important role both in mediating the perceptual experience of motion and for propagating motion information to other areas in the brain. MT provides the motion information necessary for navigating and interacting with the 3D world to numerous cortical areas and has reciprocal projections (Abe et al., 2018) to a variety of sensorimotor areas including parietal V6 and V6A (object motion recognition and control of reach-to-grasp movements (Gamberini et al., 2011; Pitzalis et al., 2013)), AIP (visuo-motor transformations for grasp (Nelissen et al., 2018)), MIP (coordination of hand movements and visual targets (Grefkes & Fink, 2005)), LIP (saccadic target selection) and frontal A4ab (motor cortex), prefrontal A8aV (frontal eye fields) and A8C (premotor).

## CONCLUSION

In conclusion, our findings provide some of the first direct evidence that both sighted and early blind individuals perceptually experience auditory motion using spatiotemporally separable filters. This contrasts with visual motion processing, which utilizes non-separable filters tuned for continuous object motion, and suggests that the statistics in sensory input constrains how object motion is encoded across different sensory modalities. Our results further suggest that, within the general constraints afforded by the statistics of the input, altered auditory experience or neural reorganization due to early blindness does refine auditory motion tunings.

